# Conserved and variable responses of the HEAT SHOCK FACTOR transcription factor family in maize and *Setaria viridis*

**DOI:** 10.1101/2022.05.19.492695

**Authors:** Zachary A. Myers, Clair M. Wootan, Zhikai Liang, Peng Zhou, Julia Englehorn, Thomas Hartwig, Nathan M. Springer

## Abstract

Responding to the environment is a core aspect of plant growth and development. Mounting an effective response is important for plants to balance growth and survival. The HEAT SHOCK FACTOR (HSF) transcription factor family is a central and required component of plant heat stress responses and acquired thermotolerance. The HSF family has dramatically expanded in plant lineages, often including a repertoire of 20 or more genes. Here we assess the composition and heat responsiveness of the HSF family in *Setaria viridis* (*Setaria*), a model C4 panicoid grass, and make targeted comparisons between the HSF families of *Setaria* and maize. Examples of both conserved and variable expression responses to a heat stress event were observed when comparing the two species. Novel and existing data on chromatin accessibility, histone modifications, and genome-wide DNA binding profiles were utilized to assess the chromatin of HSF family members with distinct responses to heat stress. We observed significant variability for both expression and chromatin state within syntenic and orthologous sets of HSFs between *Setaria* and maize, as well as between syntenic pairs of maize HSFs retained following its most recent genome duplication event. These observations collectively support a complex scenario of expansion and sub-functionalization within this transcription factor family that has significant untapped potential for better understanding the evolution of large gene families.

**Significance Statement:** A comparison of the Heat Shock Factor transcription factors in maize and *Setaria* reveals examples of consistent and variable expression responses to heat stress and provides insights into the role of chromatin in predicting expression responses.

## Introduction

Environmental perception and responsiveness are key features of plant growth and development. Understanding the intricate ways in which such capabilities have evolved and are encoded could provide avenues to develop more resilient crop varieties. A foundational component of these features lies in transcriptional gene regulation, where an interplay of transcription factors coordinates complex expression profiles. Transcription factors (TFs) are ubiquitous to all organisms, but many TF families are significantly expanded in plant lineages relative to non-plant lineages (Shiu *et al*., 2005). This expansion likely facilitates potential sub-functionalization and complex interactions.

While many plant genomes have undergone significant duplication events to ultimately produce large, highly repetitive genomes, TF families appear to have expanded to a larger extent than other genes (Shiu *et al*., 2005). Given the multiple documented whole genome duplication events in higher plants, multiple theories have gained traction to explain the higher retention rate of TFs than other genes (Panchy *et al*., 2016). In particular, the concepts of dosage balance, where maintenance of TF stoichiometry is proposed to contribute to early retention (Freeling and Thomas, 2006), and sub-functionalization, where duplication relaxes purifying selection on one of the pair of recently duplicated TFs (Ohno, 2013), have been widely accepted in a non-mutually-exclusive manner. Studies incorporating family-wide TF characterization are one avenue through which these evolutionary processes can be evaluated, and observations supporting a multitude of theories have been reported. Examples of these include the role of an expanded *MADS*-box homeotic TF family establishing the “orchid floral code (Mondragón-Palomino and Theißen, 2011),” and the impact of subfunctionalization of *AP3* TFs in establishing floral architecture in tomato (Martino *et al*., 2006).

### HSFs as a lens through which to explore TF function and evolutionary implications

Among the many expanded TF families in plants is the HEAT SHOCK FACTOR (HSF) family, a TF family that is conserved among eukaryotes. HSF families are typically small in animals and fungi, encoded by 1 and 4 genes in *S. cerevisiae* and humans, respectively (Sorger and Pelham, 1988; Nakai *et al*., 1997), but are significantly expanded in plant species, ranging from as few as 16 in *Camellia sinensis (Liu et al., 2016)*, to as many as 56 in wheat (Xue *et al*., 2014). Many aspects of HSF biology are conserved across eukaryotes, including their DNA-binding unit consisting of a multimeric complex that recognizes inverted repeats of an *NGAAN* sequence (Perisic *et al*., 1989). Plant HSFs are typified by highly conserved DNA-binding and oligomerization domains, and are sub-classified into 3 families: HSFA, HSFB, and HSFC. These families are assigned by the presence or absence of activator motifs and nuclear import and export signals, as well as the length of key feature linker sequences. Interestingly, the expansion of different HSF families in plant lineages has not been consistent, and several broad trends have been observed, including relative overexpansion of the *HSFC* family in monocots relative to eudicots and the lack of *HSFA9, HSFB3*, and *HSFB5* members in many monocots (Guo *et al*., 2016). These large and diverse families of plant HSFs are involved in many cellular processes, including mediating heat shock responses through activation of molecular chaperones known as HEAT SHOCK PROTEINs (HSPs), regulating salt and osmotic stress in transgenic Arabidopsis (Ogawa *et al*., 2007; Yokotani *et al*., 2008), and forming a potential ABA- and DREB-independent pathway to mediate drought response in transgenic Arabidopsis (Bechtold *et al*., 2013).

Prior work in plants suggests that HSFA1 is expressed and present prior to heat stress events in an inactive, sequestered state. Heat stress results in activation and nuclear localization that allows activation of target genes including the HSPs, as well as additional HSF TFs (Ohama *et al*., 2017; Kotak *et al*., 2007; Koskull-Döring *et al*., 2007). Studies in Arabidopsis have shown that the four genes encoding HSFA1s provide largely redundant function and the quadruple mutant is highly heat sensitive (Yoshida *et al*., 2011), suggesting a key role for HSFA1. There is a single HSFA2 gene in Arabidopsis. This gene is transcriptionally activated by HSFA1 and plays important roles in activating expression of downstream genes and enabling transcriptional memory in concert with HSFA3 (Nishizawa-Yokoi *et al*., 2011; Friedrich *et al*., 2021). A subset of the HSFs in Arabidopsis exhibit elevated transcript abundance following heat stress events (Swindell *et al*., 2007). Analysis of the HSF family composition and heat responsiveness have been reported for many plant species (Guo *et al*., 2016), including maize (Lin *et al*., 2011); however, the shifts in heat-responsiveness for syntenic orthologs or paralogs have not been assessed carefully in monocots. The HSFs provide a potential model gene family for considering how chromatin and cis-regulatory elements might vary to influence altered gene expression responses to an environmental cue.

In this study we utilize two C4 grasses, maize and *Setaria viridis* (*Setaria*), to assess changes in composition and heat-responsive expression of HSF genes. Maize is an agronomically important crop with significant genetic and genomic resources (Schnable *et al*., 2009; Hufford *et al*., 2021), while *Setaria* is a more recently developed model system with the benefits of a small genome (Mamidi *et al*., 2020), transformability (Thielen *et al*., 2020; Weiss *et al*., n.d.), and growth parameters well suited for greenhouse and growth chamber environments (Huang *et al*., 2016; Li and Brutnell, 2011). In contrast to dicots, there are notable shifts in the number of paralogs within many sub-clades of HSF genes. Transcriptome analyses in both species reveal many examples of conserved heat-responsiveness for HSF expression as well as examples of divergent patterns of response to heat stress. Chromatin accessibility or modification datasets were utilized to assess the properties of HSF genes with varying expression responses to heat stress. Our results highlight the complexity of varying gene content, chromatin and regulation within a family of transcription factors that play important roles in response to heat stress.

## Results

In order to facilitate targeted comparative genomic approaches between maize and *Setaria*, we identified and extracted predicted HSF protein sequences from the *Setaria viridis* A10 genome (Mamidi *et al*., 2020). In total we identified 26 putative SvHSFs, including 14 HSFAs, 7 HSFBs, and 5 HSFCs, which subdivide into a similar distribution of subfamilies as other related grasses (Figure 1, (Nagaraju *et al*., 2015; Wang *et al*.,2009). Phylogenetic analysis of *Setaria* and previously identified maize HSF TFs identified conserved subfamilies between the two species, but also uncovered a handful of compositional differences (Figure 1A). All of the HSF subfamilies in maize have putative orthologs or syntelogs in *Setaria*, with the majority of these identified as syntenic sets (Figure 1B, Table S1). Seven HSF subfamilies in maize have retained maize1:maize2 paralogs, accounting for the bulk of the difference in total HSF family size between the two species. Interestingly, it also appears that two *Setaria* subfamilies, *SvHSFA6a* and *SvHSFC2*, both have undergone tandem duplication events (Figure S2). We confirmed these HSF designations based on the presence of HSF-type DNA binding domains (DBDs) and oligomerization domains (ODs) through multiple sequence alignment against 31 previously identified maize HSFs (Figure S1). Only Sevir.1G026300 lacked significant conservation in portions of the DBD, though it did align well against the OD and subregions of the DBD, and was grouped with high confidence into a clade containing HSFC2 members in phylogenetic analysis (Figure 1A). This gene appears to be an atypical HSF TF that arose from a tandem duplication of *Sevir.1G026200;* however, it is unclear if atypical HSFs are expressed and/or functional in the few instances where they have been computationally predicted (Berz *et al*., 2019). Because of these key differences, we have assigned *Sevir.1G026300* to a separate HSF subfamily, *SvHSFC2c*,than to its paired duplicate.

**Figure 1.**
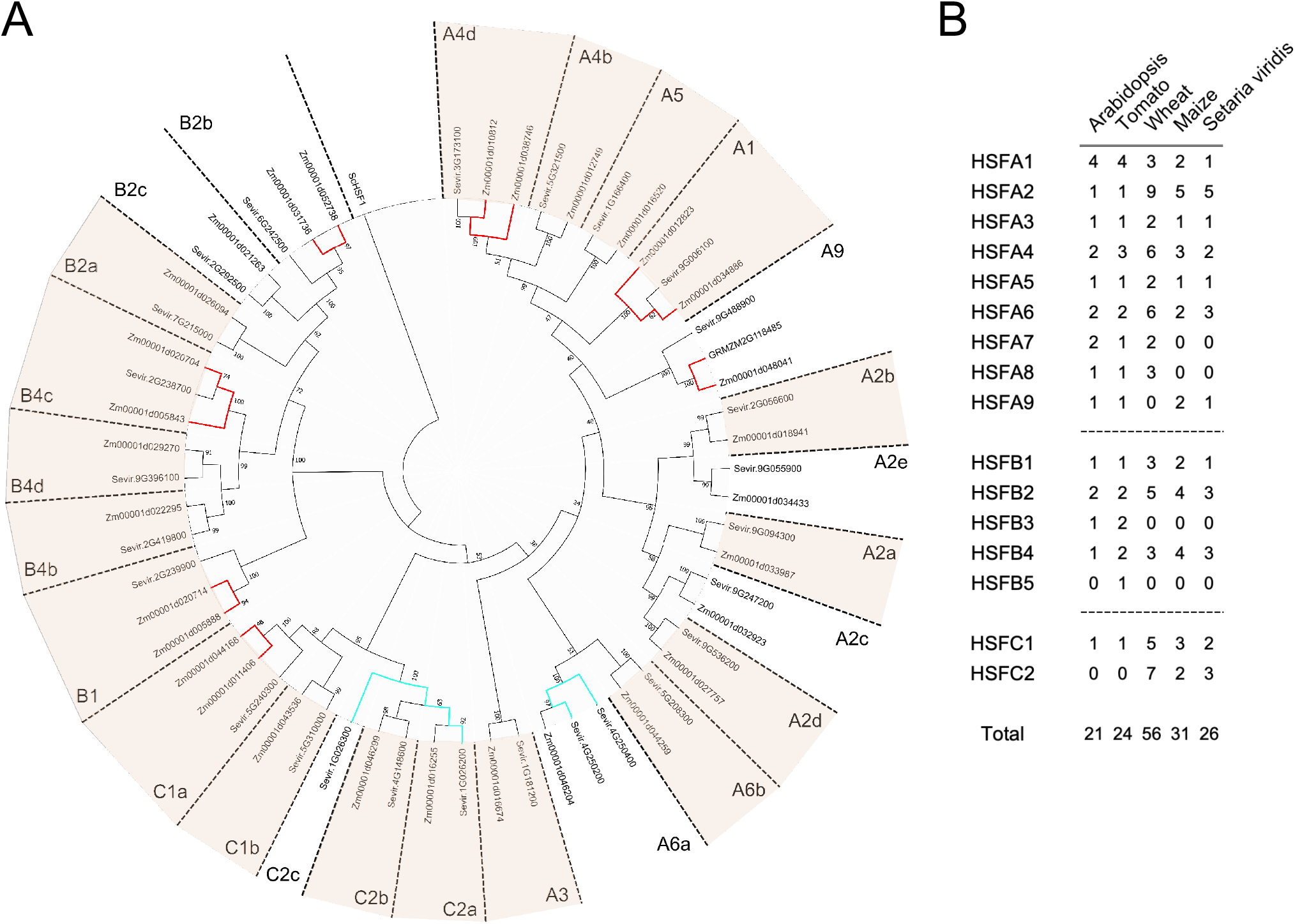
Phylogenetic relationship between the HSF families of maize and *Setaria*. (A) A neighbor-joining consensus tree was constructed to orthologous HSF subfamilies. Subfamilies are indicated on the outer ring, and were previously assigned in maize (Zhang *et al*., 2020). Putative tandem duplicates are connected in cyan, and retained maize1:maize2 paralogs are connected in red. Syntenic genes between *Setaria* and maize are shaded in tan. *Saccharomyces cerevisiae* HSF1 (ScHSF1) was included to root the tree. Numbers at branch points indicate % confidence in consensus assignment over 1000 bootstrap runs. (B) Composition of HSF subfamilies across Arabidopsis (Nover *et al*., 2001), tomato (Scharf *et al*., 2012), wheat (Xue *et al*., 2015), maize (Zhang *et al*.,2020), and *Setaria*.

### Gene expression in SvA10 in response to heat stress

With a clearer picture of the composition of the SvHSF gene family, we proceeded to assess whether orthologous HSF TFs would respond similarly to heat stress. To address this, we conducted RNA-seq experiments on control and heat stressed *Setaria* seedlings, matching growth, stress, and sampling conditions with a previously published maize dataset (Zhou *et al*., 2021). We identified 1,941 upregulated and 986 downregulated genes in response to a one-hour, 10°C increase over ambient heat stress (Figure 2A-B, Table S2). Closer examination of the differentially expressed gene sets identified a large suite of HSP chaperones that were significantly upregulated in response to heat stress (Figure 2C), and Gene Ontology (GO) enrichment of all upregulated genes identified significant overrepresentation of several heat stress responsive categories (Figure S3, Table S3). These observations collectively suggest that the applied stress effectively elicited a molecular heat response broadly similar to that observed in other species.

**Figure 2.**
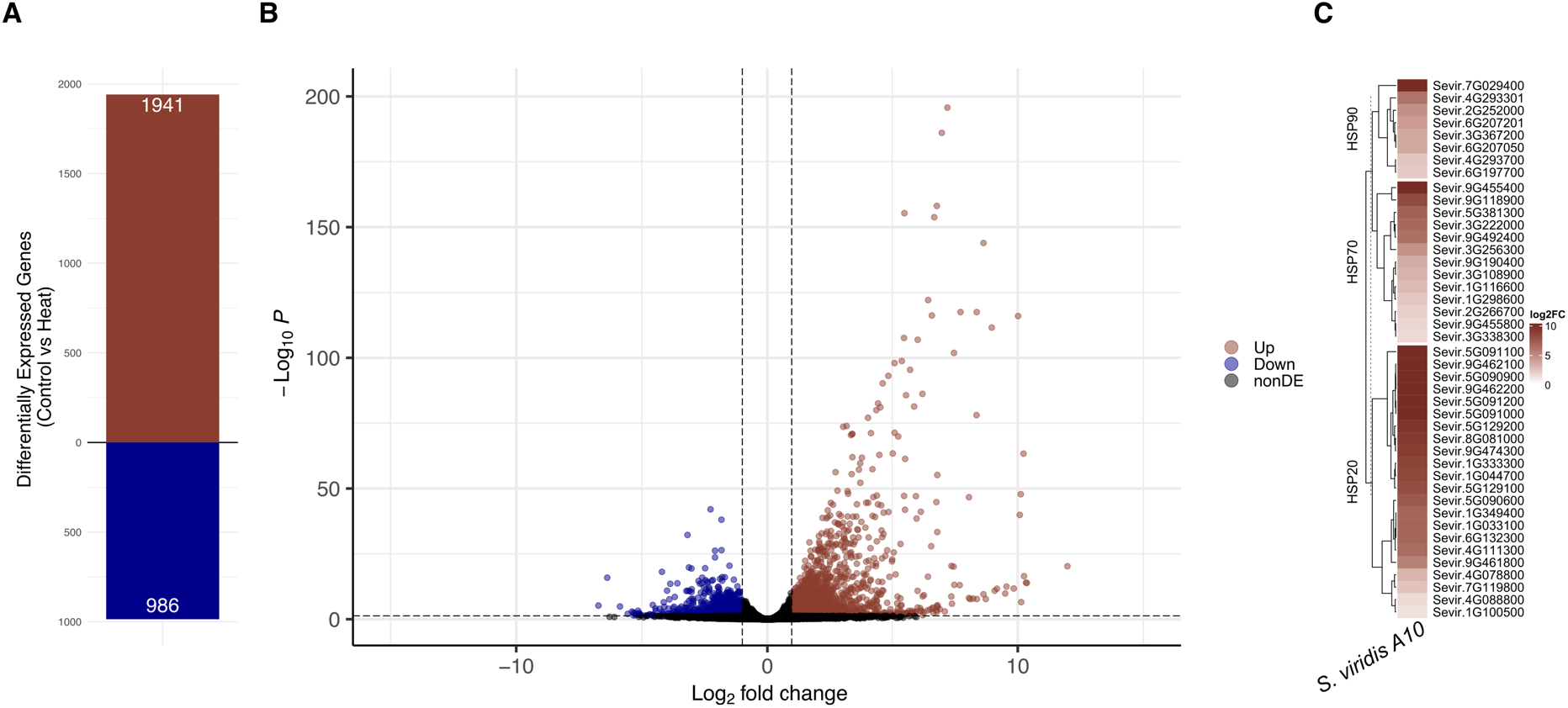
Identification of heat-responsive DEGs in *Setaria* viridis. (A) Total number of up- and down-regulated DEGs in response to a one hour, +10°C heat treatment. Cutoffs for DEG classification included an expression threshold of at least 1 count per million in one sample, a log_2_ Fold Change (heat/control) requirement of >1 (Up) or <-1 (Down), and an adjusted p-value < 0.05. (B) Volcano plot showing log_2_ fold changes and −log_10_ P values for every expressed gene. Dashed lines indicate log_2_ fold change cutoffs (vertical lines) and p value cutoffs (horizontal line). (C) Heatmap of all significantly upregulated Heat Shock Proteins (HSPs), broken down by class. Only significantly upregulated HSPs are shown here, using the same significance threshold as in (A).

### Comparison of transcriptome responses to heat in Setaria and maize

To evaluate the similarities and differences between molecular heat stress responses in *Setaria* and maize, we identified a set of 10,003 1:1 orthologs between the two species (Table S4). We found ~5-fold enrichment for orthologous genes that are up-regulated (requiring both a log_2_ fold change > 1 and an adjusted p-value < 0.05) in both species in response to heat stress (p < 3.46e-103, Figure 3A). If we reduce the stringency to require that a gene meets both criteria in one species and only has a >2-fold change or an adjusted p-value <0.05 then there are an additional 190 genes that are up-regulated in both species, suggesting that many of the non-overlapping genes are close to meeting the stringent criteria for differential expression in both species. Indeed, a visualization of the expression differences in both species finds that the genes that are differentially expressed in both species tend to exhibit higher fold-changes and that genes only significant in one of the two species often exhibit expression differences trending in the same direction in the other species (Figure 3B). The set of orthologs that are commonly up-regulated in both species exhibit a significant enrichment for heat-responsive GO terms in this overlapping set (Figure S3, Table S3). Among the genes significantly upregulated in response to heat in both species, members of the HSP class of proteins accounted for nearly 10% of all shared significantly upregulated orthologs, and were generally upregulated to a similar degree in both species (Figure 3C). These results suggest overall similarities in the gene expression responses to heat stress in maize and *Setaria*.

**Figure 3.**
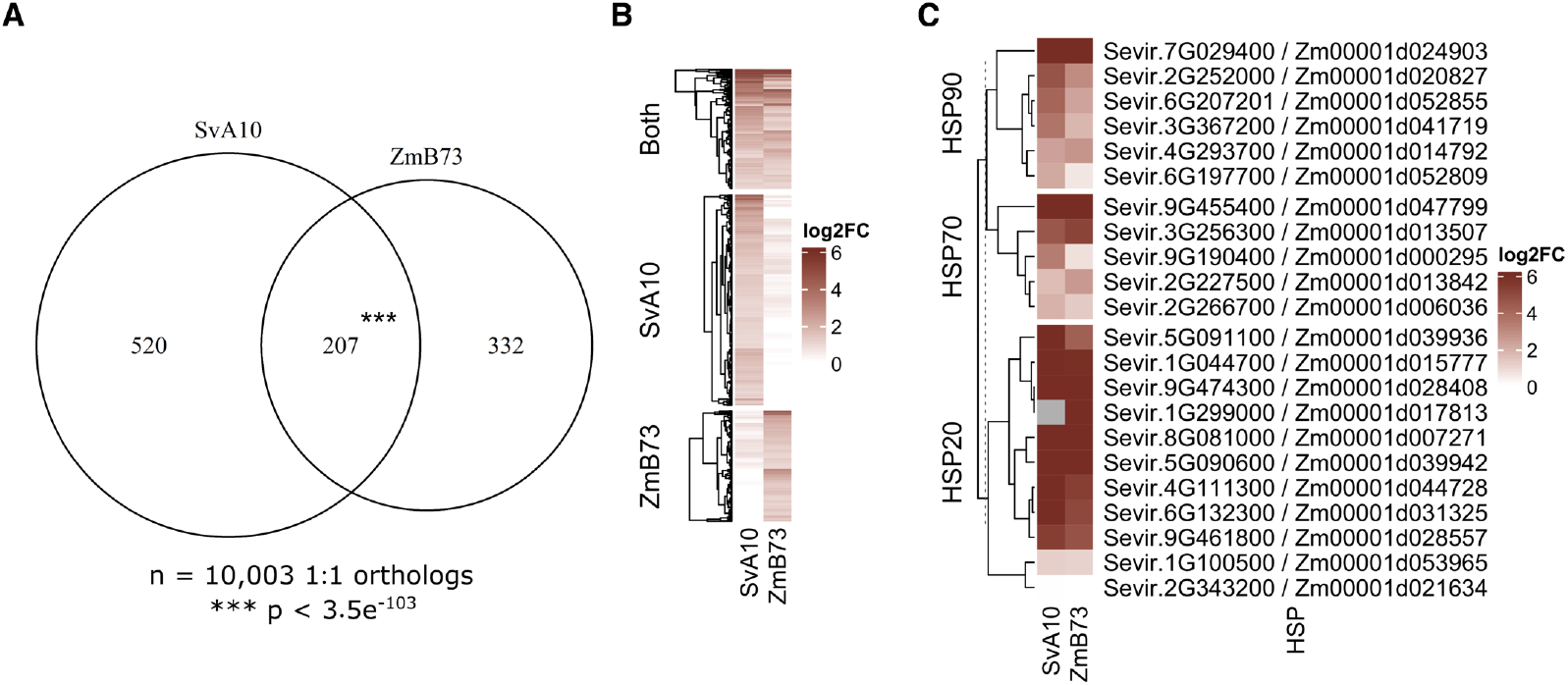
Comparison of heat responsive DEG profiles of maize and *Setaria*. (A) Overlap between 1:1 orthologs that are significantly upregulated in at least one of the two species. Asterisks indicate a significant enrichment over what could be expected by chance, calculated through a hypergeometric enrichment test. (B) Heatmap of log_2_ fold change (heat / control) values for all genes contained in (A). (C) Heatmap of all 1:1 HSP orthologs that are significantly upregulated in at least one of the two species, broken down by class.

### Comparison of heat stress responses for Setaria and maize HSFs

We examined expression levels of the HSF TFs in order to identify patterns of HSF responsiveness between the two species and differences within individual subfamilies in each species (Figure 4, S4). Members of seven HSF subfamilies were significantly upregulated in both maize and *Setaria:* A2a, A2c, A6a, B1, B2b, B2c, and C2a. These include most of the genes with the largest expression responses to heat stress. Five subfamilies exhibited *Setaria*-specific responses to heat based on significant expression responses to a one-hour heat stress event (Figure 4). Several of these examples show evidence for a heat response in maize but it was not statistically significant (Figure S6). A prior study monitoring heat responses in maize (Zhou *et al*., 2021) generated unreplicated time-course data for responses to heat stress and included a 30-minute time point. Three of the maize genes (*HSFA2b, HSFA2d*, and *HSFB2a*) from sub-families classified as having *Setaria*-specific responses exhibit striking up-regulation in response to heat stress at 30 minutes, but the response has largely subsided by 1 hour and the genes are not detected as significant at 1 hour. These may represent different dynamics of the timing of response rather than true species-specific differences in responsiveness. One subfamily exhibited a maize-specific response based on differences in significant expression responses (Figure 4). Examining the two subfamilies containing *Setaria* tandem duplicates, the *HSFA6a*subfamily was among the most heat-responsive of all HSFs in both species, while the *SvHSFC2a/SvHSFC2c* pair clearly diverged, with the *HSFC2a* subfamily exhibiting conserved heat responsiveness in both *Setaria* and maize, but no detectable expression of *SvHSFC2c* (Figure 4, S4).

**Figure 4.**
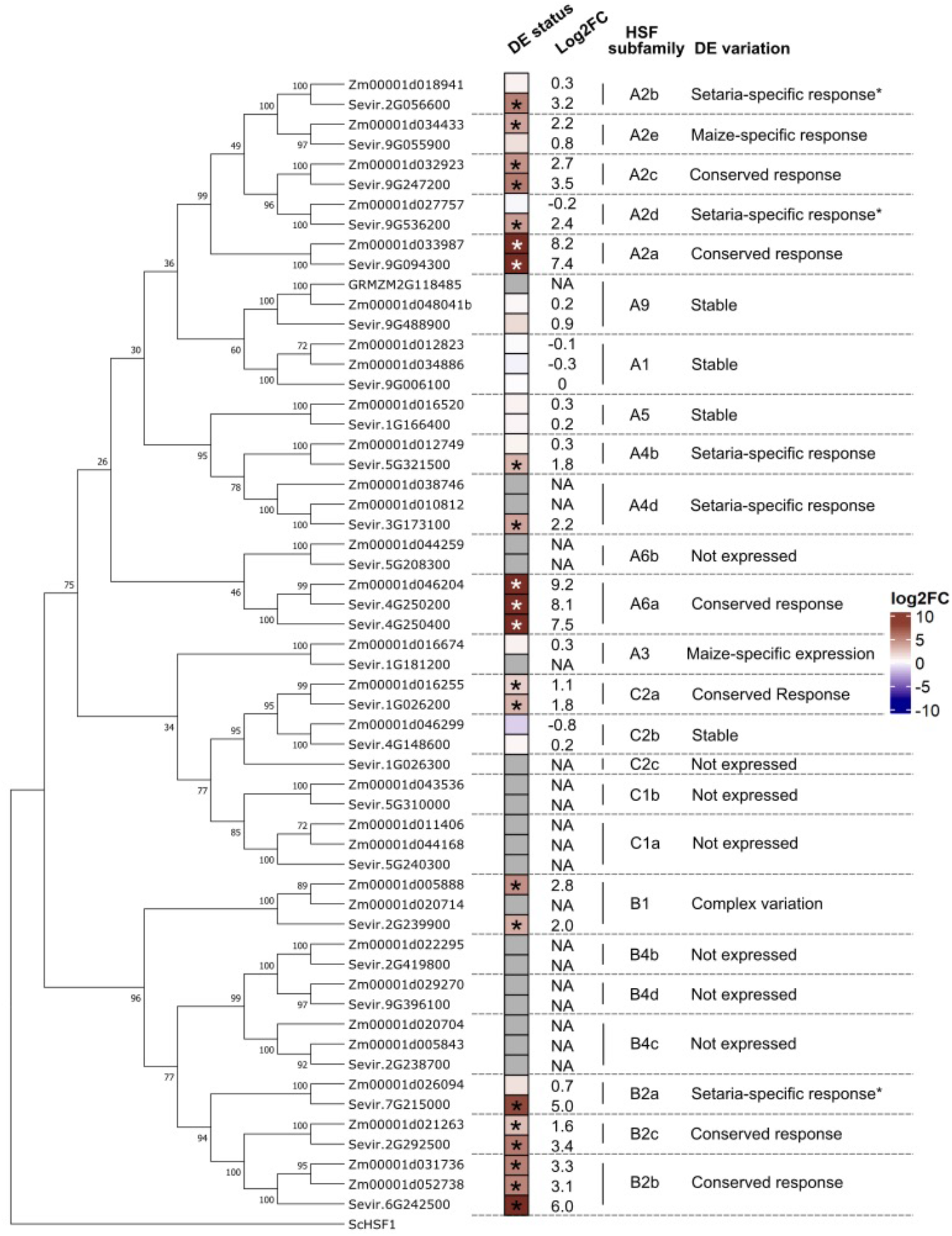
Subfamily-based comparison of HSF heat responsiveness between maize and *Setaria*. Phylogenetic tree is reformatted but otherwise identical to that presented in Figure 1. DE status is presented as a heatmap of log_2_ fold changes, with not expressed HSF (< 1 CPM) colored gray with actual log_2_ fold change values are included in the adjacent column. HSF subfamilies are indicated, as well as a brief explanation of the observed variation. Asterisks indicate three subfamilies identified as *Setaria*-specific responsive where prior time course evidence suggests a missed heat responsive pattern (e.g., these maize HSFs may have responded and returned to pre-stress levels by the time tissue was collected).

While there are no maize tandem duplicate HSFs, we did more closely examine the previously identified maize1:maize2 paralogs through the lens of a previously published maize gene expression tissue atlas (Stelpflug *et al*., 2016; Walley *et al*., 2016; Zhou *et al*., 2020; Chen *et al*., 2014; Yi *et al*., 2019; Zhou *et al*., 2019). We extracted normalized expression levels for six paralog pairs across 247 RNA-seq experiments and performed hierarchical clustering (Figure S5). While several of these pairs were not expressed in the leaf tissue heat stress datasets used here, we were able to identify tissues and/or conditions where each was expressed. In some cases (HSFA1, HSFB1) there is evidence for higher expression of one of the two paralogs (Figure S5). In other cases (such as B4c), the two duplicate genes have highly similar patterns of expression. There are also examples such as A4d and C1a in which the tissue-specific patterns of expression of the retained duplicates have diverged (Figure S4).

### Chromatin and epigenomic feature analysis of HSFs

The HSF gene family provides an example of a gene family with variable regulation for different family members. We were interested in using new and previously generated chromatin and epigenomic datasets to compare the chromatin at genes with variable responses to heat stress. Each of the HSF genes was classified into four categories based on expression levels in control compared to heat samples: (1) Increased (expressed in control, significantly up in heat), (2) Stable (expressed in both conditions), (3) Activated (not expressed in control, significantly up in heat), and (4) not expressed (Figure 5A, S5). The groups were fairly evenly populated in *Setaria*, with 5 ‘Stable’ HSFs, 7 ‘Increased’ HSFs, 6 ‘Activated’ HSFs, and 8 ‘not expressed’ HSFs. In maize, however, there were only 2 ‘Activated’ HSFs, as well as 7 ‘Increased’, 10 ‘Stable’, and 11 ‘not expressed’ HSFs.

**Figure 5.**
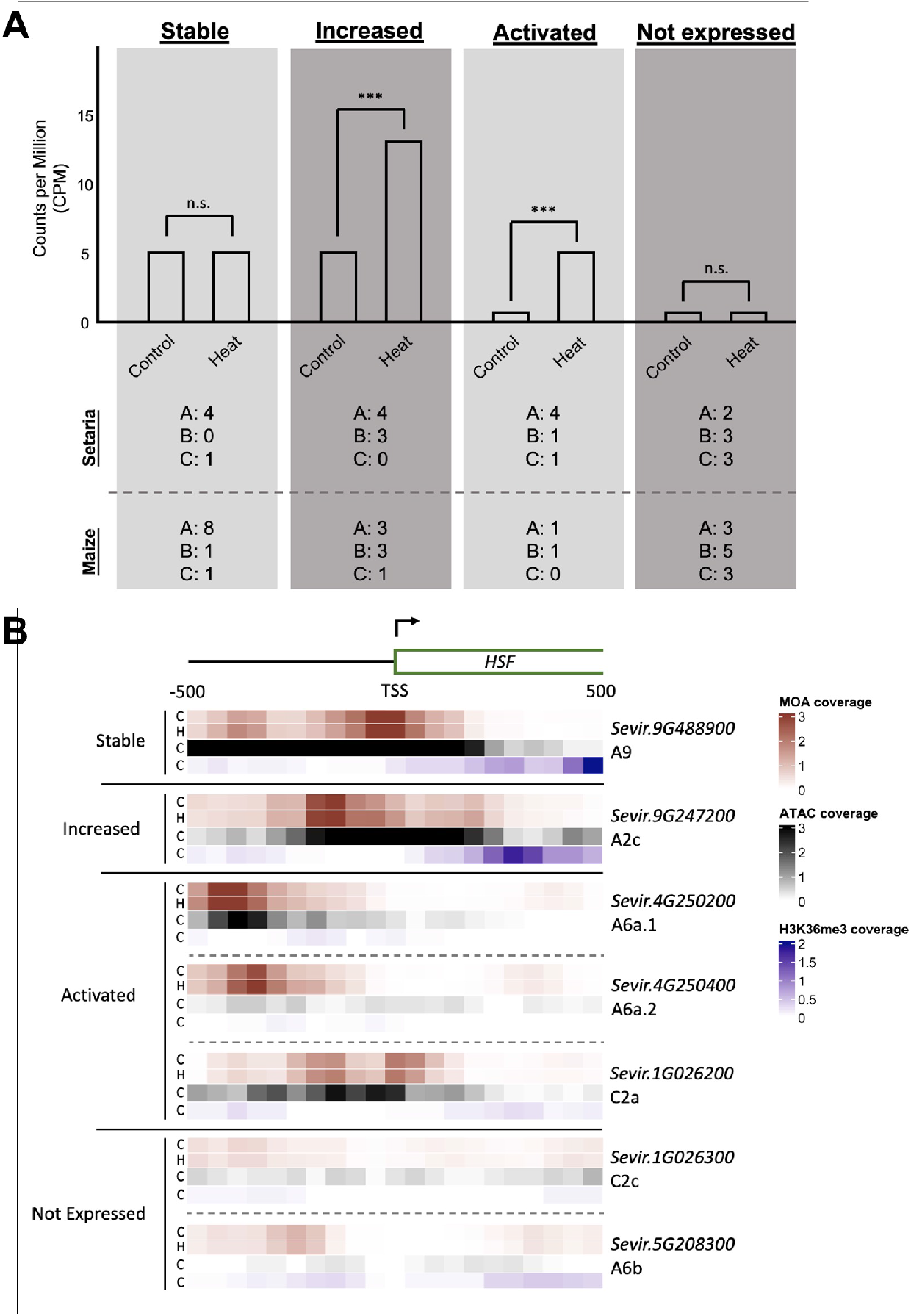
Expression and chromatin modification analysis of the *Setaria* and maize HSFs. (A) Schematic representation of the four expression categories to which each HSF was assigned. Asterisks indicate significant DEG calls between control and heat conditions; n.s., not significant. The total number of HSFs in each subfamily (HSFA, HSFB, and HSFC) assigned to each expression category is listed separately for *Setaria* and maize below the schematic. (B) Chromatin accessibility and modification datasets were used to generate a heatmap of coverage over a 1kb region centered on the TSS of representative members for each expression category. A generic gene model depicting the genomic region is included at the top of the panel. One or more representatives of each expression category are included and indicated by group on the left side of the heatmap, with specific gene IDs and subfamily assignments on the right. The “C” and “H” indicate Control and Heat-stressed samples, respectively.

We generated global scale TF footprinting datasets in both control and heat-stressed (1h, +10C) conditions through MOA-seq (MNase-defined cistrome Occupancy Analyses-sequencing, (Savadel *et al*., 2021)), and leveraged previously published chromatin accessibility (ATAC-seq, Assay for Transposase-Accessible Chromatin-sequencing) and H3K36 trimethylation Chromatin Immunoprecipitation-sequencing levels (H3K36me3 ChIP-seq, (Lu *et al*., 2019)). While we generated MOA-seq data for both control and heat-stressed tissue samples, the prior chromatin datasets were all generated only for plants grown in control conditions. We initially focused on the region surrounding the transcriptional start site (TSS) of the *Setaria* HSFs, examining the patterns of TF binding, chromatin accessibility, and H3K36 trimethylation levels (Figure 5B, S6). While there was certainly variation within groups of genes with similar expression responses, a handful of trends stood out when comparing the expression categories across these genomics datasets. In categories expressed in ambient conditions (‘Stable’ and ‘Increased’), we tended to see large open chromatin regions and smaller regions of MOA signal peaks. Genes that are expressed in ambient conditions also tend to have high levels of H3K36me3 in the region immediately downstream of the TSS. HSFs in the ‘Activated’ category, which are not expressed at ambient temperature, tended to have some level of chromatin accessibility and typically still had consistent MOA signal peaks, suggesting the presence of accessible chromatin prior to activation of expression. Finally, HSFs in the ‘not expressed’ category tend to lack evidence for accessible chromatin and H3K36me3. We did not find evidence for significant differences in MOA signal between control and heat conditions, even at HSF genes with substantial increases in expression in heat.

A similar set of analyses were performed for the maize HSFs with one notable difference in sampling: the MOA-seq data was generated for leaf tissue samples after a 4 hour +10°C stress treatment rather than a one-hour treatment. Focusing on the region around each HSF TSS and faceting by expression category, we saw many of the same trends as observed in *Setaria*, with a few key differences (Figure 6, S7). Specifically, while H3K36me3 levels seemed relatively similar across both species when comparing individual expression categories, chromatin accessibility was observed in more ‘not expressed’ HSFs in maize than in *Setaria*, and the size of accessible chromatin regions in maize appeared more restricted. Further, while the patterns of MOA signal seemed relatively similar between the two species across expression categories, there were several instances of noticeably increased MOA signal in response to heat stress in maize but not *Setaria*. One possible explanation for the evidence for altered MOA-seq peaks in maize but not *Setaria* could be the later sampling time in maize compared to *Setaria*.

**Figure 6.**
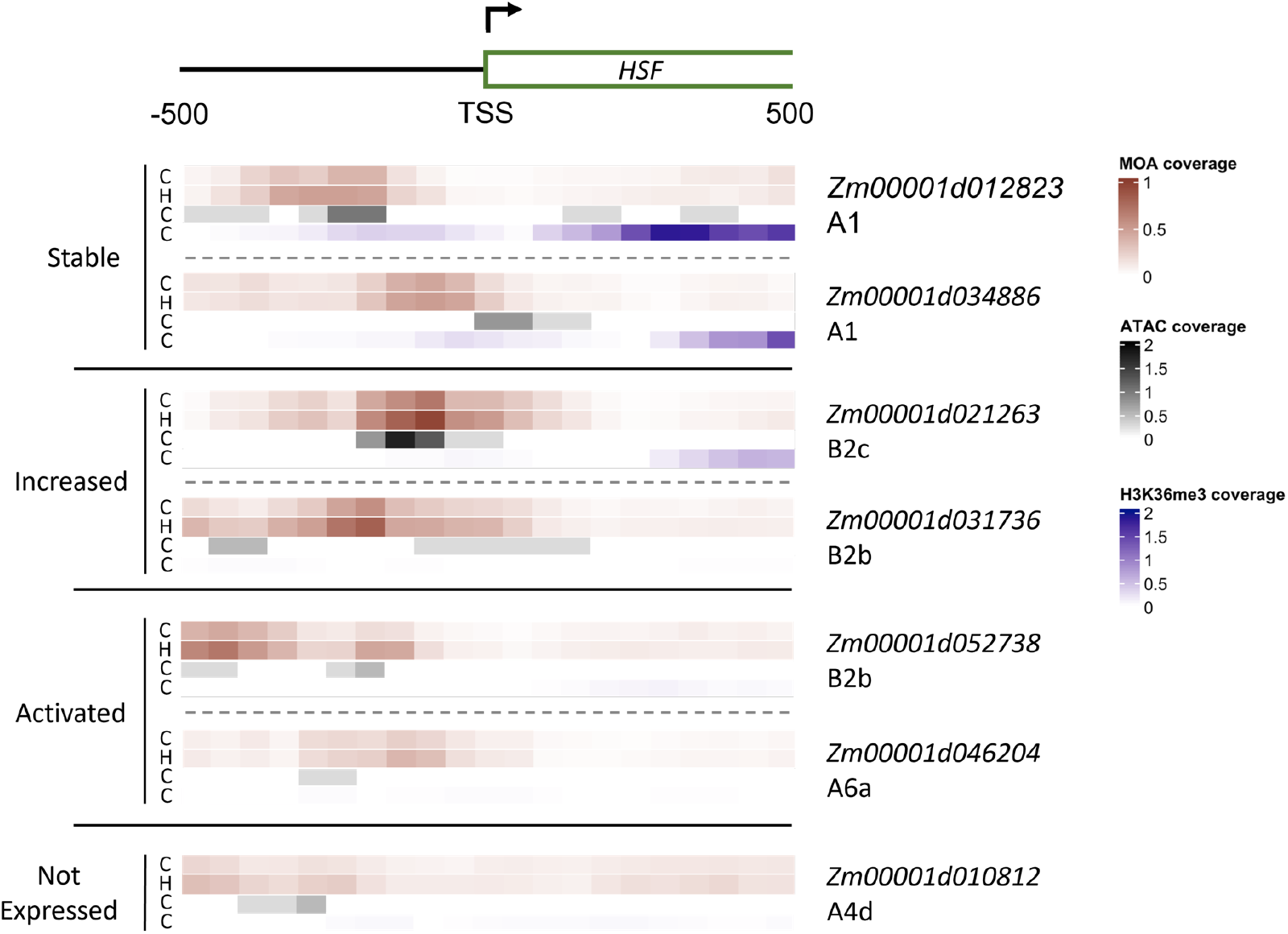
Chromatin accessibility and modification analysis of maize HSFs. Chromatin accessibility and modification datasets were used to generate a heatmap of coverage over a 1kb region centered on the TSS of representative members for each expression category. A generic gene model depicting the genomic region is included at the top of the panel. One or more representatives of each expression category are included and indicated by group on the left side of the heatmap, with specific gene IDs and subfamily assignments on the right. The “C” and “H” indicate Control and Heat-stressed samples, respectively.

### Comparisons of chromatin properties at paralogs and orthologs/syntelogs

In addition to exploring expression category-wide trends, we also made targeted comparisons of paralogs and orthologs. There are two pairs of *Setaria* tandem duplicates that are present in *Setaria* but not maize. These two pairs of tandemly duplicated genes provide an opportunity to consider the patterns of chromatin at duplicated genes. When examining the *HSFC2a/HSFC2c* pair, which split into different expression categories (‘not expressed’ and ‘Activated’), we observed a clear reduction in both MOA and ATAC signal in *HSFC2c* compared to *HSFC2a* (Figure 5B, S8). Taken alongside the complete lack of expression of *HSFC2c* in our datasets, it seems that *HSFC2c* may be in the process of pseudogenization. The tandem duplicate pair of *HSFA6a* members were both assigned to the ‘Activated’ category and exhibit similar responses. Both genes lack evidence for H3K36me3, as expected due to the lack of expression in control conditions. While the promoters of these two HSFA6a members have nearly completely diverged, they both have notable MOA-seq signal, though the relative location is different (Figure S8). *Sevir.4G250200* has a strong ATAC peak that overlaps a MOA-seq peak but there is no evidence for an ATAC-seq peak at *Sevir.4G250400* (Figure 5B, S8).

While there are no examples of HSF tandem duplications in maize, there are several examples of retained duplicates from a whole genome duplication event (termed maize1:maize2). Four maize1:maize2 orthologous HSF pairs were assigned to the same expression category (A1, stable; A4d, B4c, and C1a, not expressed), while two pairs were assigned to different expression categories (B1, ‘Increased’ and ‘not expressed’; B2b, ‘Increased’ and ‘Activated’). The chromatin and epigenetic landscape around the TSSs of the stably expressed HSFA1 pair in maize was remarkably similar, though small (~100bp) shifts in the relative positions of MOA and ATAC signal were observed (Figure 6). The ‘Increased’ and ‘Activated’ pair of HSFB2b members in maize also displayed a similar but shifted landscape near their respective TSSs, though perhaps with a more noticeable increase in MOA signal in the heat-stressed samples compared to control (Figure 6).

Expression categorization of individual HSF subfamilies across species was not particularly correlated, with only 10 of the 22 subfamilies assignments consistent between *Setaria* and maize. The HSFA6a subfamily was one where all members were assigned to the same expression category (‘Activated’), and a comparison of the chromatin and epigenetic landscape around their respective TSSs largely mirrored the differences observed between the two species at large - all three members were void of H3K36 trimethylation, had MOA signal upstream of the TSS, and while all had some level of chromatin accessibility, it was markedly higher in one of the *Setaria* HSFA6a members (*Sevir.4G250200*) compared to either of the other two (*Sevir.4G250400* and *Zm00001d046204)*. Comparison of the *HSFB2b* subfamily, which is encoded by a retained m1:m2 paralog in maize and a single locus in *Setaria*, identified similar patterns of chromatin accessibility and TF binding (Figures 6, S6) despite assignment to different expression categories (‘Increased’ and ‘Activated’ in the two maize copies, ‘Increased’ in *Setaria*). While ATAC signal was relatively low around the maize *HSFB2b* TSS, it was overlapping with or near MOA signal - particularly in heat stress conditions (Figure 6). In *Setaria HSFB2b*, accessibility was markedly increased and was positioned over a region of increased TF binding in response to heat stress (Figure S6). The *HSFB1* subfamily is also encoded by a retained m1:m2 paralog in maize and a single locus in *Setaria* (Figure 1). This subfamily was again assigned to different expression categories, with an ‘Increased’ and ‘not expressed’ HSF in maize and an ‘Increased’ HSF in *Setaria* (Figure S4). Interestingly, MOA signal was similar in all three *HSFB1* members, with a notable peak upstream of the TSS; however, the two ‘Increased’ HSFs have open chromatin regions, while the ‘not expressed’ HSF is not accessible near the MOA signal region (Figures S6, S7). The lack of open and quickly accessible chromatin, such as that seen in the two ‘Increased’ HSFB1 members, might contribute to the lack of heat-responsive expression.

## Discussion

### Differential HSF family expansion in monocots and dicots

The HSF family has been broadly identified in many plant species, and as described above, a number of trends have been described when comparing monocot and dicot lineages. One interesting aspect that has received relatively little attention is the functional consequences of expansion and diversification of the *HSFA1* and *HSFA2* families between the two lineages. The function and relative contribution to heat stress responses of these two subfamilies have been dissected in Arabidopsis and tomato (*Solanum lycopersicum*), where a constitutively expressed and inactive HSFA1 is functionally activated in response to heat stress and subsequently drives expression of many heat-responsive genes, including *HSFA2* (El-Shershaby *et al*., 2019; Yoshida *et al*., 2011; Li *et al*., 2010). Interestingly, while both Arabidopsis and tomato encode 4 *HSFA1* genes, only one is necessary for driving heat stress responses in tomato, while at least 3 of the Arabidopsis *HSFA1* genes redundantly regulate the same general heat stress response. A recent report suggested that the non-essential tomato *HSFA1* members have functionally diverged, each acquiring novel tissue-specific expression and preferential activity on specific genes as a result of a mutation within the otherwise-conserved DNA binding domain (El-Shershaby *et al*., 2019). As a final piece of the puzzle to consider, tomato *HSFA1* and *HSFA2* have been suggested to form hetero-oligomeric superactivator complexes on key heat-responsive promoters, where increased transcriptional activity of target genes required a functional DNA binding domain in one of the members of the oligomer and different combinations of C-terminal activation domains (Chan-Schaminet *et al*., 2009). These pieces of data suggest a highly complex and redundant system of responding to heat stress, where individual HSF properties are often less significant than the sum of the parts within the TF complex. Given this, the consequences of family expansion and diversification will often need to be teased apart on a case-by-case basis.

In the cases of *Setaria* and maize, composition and responsiveness of the HSFA1 and HSFA2 subfamilies were markedly different than those described in Arabidopsis and tomato. Broadly speaking, the sizes of these subfamilies are swapped between the two groups of species, with *Setaria* and maize encoding fewer *HSFA1*s (1 and 2, respectively) and more *HSFA2s* (5 in each). Further, the regulation of *HSFA2* subfamilies seemed markedly different, where all but one of the ten *HSFA2* TFs in *Setaria* and maize were expressed in control conditions and seven of the 10 were heat responsive (Figs 4, S4). One possible explanation for the relative heat tolerance of maize and *Setaria* compared to that of Arabidopsis and tomato could be that these two grass species are expressed constitutively, providing some basal level of heat tolerance.

### Broad strokes of similarities and differences in HSFs between Setaria and maize

A comparison of the full suite of HSF genes in maize and *Setaria* revealed many similarities and several key differences. We found many of the previously reported trends in maize HSF family composition (Lin *et al*., 2011; Zhang *et al*., 2020) to hold true in *Setaria*, including an expanded *HSFC* subfamily (relative to dicots) and the absence of the *HSFB3* and *HSFB5* subfamilies. Compositional differences between the HSF families of *Setaria* and maize are fully explained by either retained maize1:maize2 paralogs or tandem duplications in *Setaria*. The expression responses of HSFs to heat stress were more varied between the two species. While many HSFs have similar or marginally different responses, there are examples such as *HSFA4d* and *HSFB2a* that exhibit heat-responsive expression in *Setaria* alone. Relatively little is known about the function of these HSF subfamilies. The *HSFA4d* subfamily has been explored somewhat in rice, where overexpression of *OsHSFA4d* exhibited increased heat tolerance in transgenic rice (Yamanouchi *et al*., 2002), and overexpression of the related wheat HSFA4a led to improved Cadmium tolerance in transgenic rice (Shim *et al*., 2009). The *HSFB2* subfamily has been observed as heat stress induced in pigeonpea (*Cajanus cajan*) and implicated in defense responses against Botrytis in cucumber (*Cucumis sativus*) through induction of pathogenesis-related defense proteins in response to heat stress (Kharisma *et al*.,2022; Ramakrishna *et al*., 2022). Maize *HSFB2a* was stably expressed in the one hour heat stress data used here, but a previous study suggested that it was transiently induced at thirty minutes post heat stress before returning to its pre-stress expression level (Zhou *et al*., 2021), suggesting a potentially different temporal response between the two species.

### HFSA6 subfamily diversification in the grasses

The *HSFA6* family has been cursorily examined in wheat, where it was found to be heat induced and able to drive enhanced heat responsiveness when ectopically induced alongside a heat stress (Xue *et al*., 2015). With this in mind, it is interesting to note that *Setaria* has retained a tandemly duplicated copy of HSFA6a that shows a similar heat responsive profile. Across plant lineages, the HSFA6 family appears to typically be encoded by 2 or 3 members (Guo *et al*., 2016), though there are no identified members in apple or carrot (Giorno *et al*., 2012; Huang *et al*., 2015), and as many as 6 members in wheat (Xue *et al*., 2014). Despite the significant level of induction in response to heat observed in maize and *Setaria*, we can find no reports of Arabidopsis HSFA6a being heat responsive, either in the literature or through the Arabidopsis eFP browser (Winter *et al*.,2007). Further, we discovered that the tandem duplication event in *Setaria* appeared to also encompass a putative HSF TFBS approximately 200bp upstream of each HSFA6a coding sequence, while the sequences surrounding these HSF TFBSs had nearly completely diverged. When examining the maize HSFA6a gene, we noticed an HSF TFBS positioned several hundred base pairs upstream of the HSFA6a coding sequence (Figure S8 in *Setaria*, not shown in maize). The presence of this motif in multiple species, as well as its conservation following tandem duplication in *Setaria*, suggests that this HSF subfamily could itself be driven by another HSF.

### Use of chromatin properties to predict expression responses in a gene family

The expansion of TF families has provided opportunities for specialization and sub-functionalization of individual family members. Divergent patterns of gene expression for members of gene families can be one mechanism of sub-functionalization. HSFs often exhibit responsiveness to heat stress events, but this is not necessarily observed for all members of the gene family. In many species, researchers have used gene specific approaches (such as quantitative real-time PCR) or transcriptome profiling following a heat stress to monitor which HSF genes exhibit response to heat stress. We were interested in assessing whether chromatin accessibility or chromatin modifications could be used to predict the heat responsive expression for different members of a gene family.

Prior studies (Lu *et al*., 2019; Ricci *et al*., 2019) have documented genome-wide patterns for a number of histone modifications and chromatin accessibility in maize seedling leaf tissue. We examined these features to see whether they might facilitate predicting HSF heat responsiveness. Some chromatin features, such as H3K36 trimethylation, were highly associated with genes that are expressed in ambient conditions, including the ‘Stable’ and ‘Increased’ HSFs in both species. In addition, the majority of the genes that exhibit expression in ambient or heat stress conditions contain accessible regions near the TSS. However, we did not find evidence for specific chromatin features that were found in the HSFs that are heat responsive, suggesting limited potential to use these chromatin properties to predict responsiveness. Chromatin accessibility was often found at the ‘Activated’ HSFs even though these are not expressed in ambient conditions. This may reflect the fact that many of these HSFs are rapidly (<30minutes) induced in response to a heat stress event and chromatin accessibility is necessary to allow this rapid activation.

We also generated MOA-seq data for both ambient and heat stress conditions to assess potential changes in chromatin accessibility and TF binding at genes with expression that is increased or activated in response to heat stress. For reasons beyond the scope of this study, the MOA-seq data was generated at a 1-hour time stress for *Setaria* but a 4-hour heat stress in maize. We found quite limited changes in MOA-seq profiles in *Setaria* control and heat stressed plants despite highly distinct transcript levels for many HSF genes. These genes tended to already have MOA-seq peaks in control samples suggesting potential TF binding even prior to activation. One possible explanation is that these HSFs are being actively repressed, allowing for a quick gene expression change in response to de-repression at these positions, similar to the regulation of auxin responsive genes following the de-repression of Aux/IAA repressors (reviewed in (Teale *et al*., 2006; Chapman and Estelle, 2009)). In maize plants subjected to a 4-hour heat stress we found more examples of increased MOA-seq peaks at some heat responsive genes. In many cases the maize genes have a MOA-seq peak in control conditions but the strength of the MOA-seq signal is increased in the 4-hour heat stress sample. This could reflect increased TF binding and occupancy following heat stress. Prior studies that have compared chromatin accessibility in control and stress conditions have reported similar findings of limited numbers of novel accessible regions but more examples of quantitative shifts in chromatin accessibility (Reynoso *et al*., 2021; Raxwal *et al*., 2020; Lee and Bailey-Serres, 2019; Maher *et al*., 2018).

While our analyses of chromatin properties were focused on the HSF gene family, there are further opportunities to compare heat responsive expression and chromatin in the full set of heat responsive genes in maize and *Setaria*. Understanding the potential role of chromatin accessibility and TF footprinting to document *cis*-regulatory elements and predict expression responses will provide a roadmap for understanding the evolution of gene expression responses to abiotic stress events.

## Materials and Methods

### Plant materials and growth conditions, sampling conditions

*Setaria* plants were grown in growth chambers at 30°C/20°C under 12-h light/12-h dark cycles for either twelve (RNAseq samples) or twenty (MOA-seq samples) days before harvesting leaves. Seeds were sown directly onto wet soil (PGX Gro-Mix) and bottom watered every other day until just prior to treatment or collection. All heat stress treatments were conducted approximately 3 hours after lights were turned on, and plants were moved into either a 40°C incubator or a 30°C incubator for one hour. After treatment, leaves were collected in paper envelopes and snap frozen in liquid nitrogen. Leaf 3 of twelve day old samples used in RNA-seq analyses, while both L3 and L4 were harvested for twenty day old MOA-seq samples to increase total biomass collected.

For the maize MOA-seq experiments reported here, maize B73 seedlings were grown for 3 weeks in growth chambers at 30°C/20°C under 12-h light/12-h dark cycles. Approximately 3 hours after lights were turned on, plants were exposed to either a 40°C heat stress or left at 30°C for 4 hours. Leaves 3 and 4 were collected in paper envelopes and snap frozen in liquid nitrogen, collecting a total of ~10g of fresh tissue.

### RNA isolation, library prep and sequencing, DEG calling

RNA was isolated from snap-frozen and ground leaf tissue using the Qiagen RNeasy kit, following manufacturer’s instructions without modification. Sequencing libraries were prepared using the standard TruSeq Stranded mRNA library protocol and sequenced on a NovaSeq S4 flow cell as 150-bp paired-end reads, producing at least 20 million reads for each sample. Both library construction and sequencing were carried out at the University of Minnesota Genomics Center. Sequencing reads were then processed through the nextflow-core RNA-seq pipeline (Ewels *et al*., 2020) for initial quality control and raw read counting. In short, reads were trimmed using Trim Galore!and aligned to the *Setaria* viridis reference genome (version 2.1, (Mamidi *et al*., 2020)) using the variant-aware aligner Hisat2 (Kim *et al*., 2015). Uniquely aligned reads were then counted per feature by featureCounts (Liao *et al*., 2014). Differentially expressed genes were called through the DESeq2 package in R (Love *et al*., 2014), masking genes for expression by requiring genes to have > 1 CPM in at least one sequenced library.

### Data visualization

Most figures were generated in R v4.1.1 (R Core Team (2021)), making heavy use of the ggplot2 and patchwork packages. Volcano plots were generated through the EnhancedVolcano package (https://github.com/kevinblighe/EnhancedVolcano). Heatmaps were generated through the ComplexHeatmap package (https://jokergoo.github.io/ComplexHeatmap-reference/book/). Figures were composed in Inkscape (https://inkscape.org/) and GIMP (https://www.gimp.org/).

### MOA-seq data generation and processing

*Setaria* and maize MOA-seq libraries were constructed using a previously described protocol (Savadel *et al*., 2021). Indexes of reference genomes were built using STAR (v2.7.9a) (Dobin *et al*., 2013) and raw MOA-seq data reads were preprocessed using SeqPurge (v2019_09) (Sturm *et al*., 2016). Paired reads with overlapping regions were merged into single-end reads using bbmerge (version 38.18) (Bushnell *et al*., 2017). Processed reads were aligned to the relative *Setaria* or maize reference genome using STAR. Alignment files were converted into bam formats using SAMtools (v1.9) (Li *et al*., 2009). Alignment fragments with less than 81 bp and MAPQ as 255 were kept for analysis.

### Phylogenetic tree construction, MSAs

Phylogenetic trees were generated through MEGA (Kumar *et al*., 2018), using full length amino acid sequences for each primary gene model for both maize and *Setaria* HSFs. *Setaria* HSFs were identified and confirmed through three approaches: (1) extraction of genes assigned to the HSF-type PFAM ID PF00447 in the S. viridis A10 v2.1 and S. viridis ME034v genome assemblies (Mamidi *et al*., 2020; Thielen *et al*., 2020); (2) comprehensive searching through the HSF-centered HEATSTER database (Scharf *et al*.,2012); and (3) querying the S viridis A10 v2.1 genome through BLAST with all previously identified maize HSFs. Neighbor-Joining trees were constructed using 1,000 bootstrap iterations to produce the final consensus trees presented here (Saitou and Nei, 1987). Multiple sequence alignments were generated in Geneious Prime (v11.0.11) using the MUSCLE alignment algorithm with default parameters and up to 1,000 bootstrap iterations (Edgar, 2004). As presented in the main text, full amino acid sequence MSAs were trimmed down to the DNA binding domain region and/or oligomerization domain region in some instances.

### Identification of syntenic genes and paralogs

Syntenic genes and sets were pulled from an extension of a previous analysis (Schnable *et al*., 2011; Zhang *et al*., 2017)). Identification of all 1:1 orthologs between maize and *Setaria* was performed through Orthofinder v2.5.2 (Emms and Kelly, 2019), using primary amino acid sequences for the Maize B73v4 assembly (Jiao *et al*., 2017) and the *Setaria* A10 v2.1 assembly (Mamidi *et al*., 2020). Orthofinder was run with default parameters, and 1:1 orthogroups were extracted to construct a final 1:1 ortholog table.

### Data availability and use of previously published datasets

Previously published ATAC and H3K36 trimethylation ChIP datasets (Lu *et al*., 2019) were retrieved from the NCBI Sequence Read Archive, accession number GSE128434, through use of the sra-toolkit (https://github.com/ncbi/sra-tools). Raw reads were adapter trimmed with Trimmomatic (Bolger *et al*., 2014), quality controlled with FastQC (https://www.bioinformatics.babraham.ac.uk/projects/fastqc/), and aligned to either the Maize B73v4 or *Setaria* A10v2 genome with bowtie2 (Langmead and Salzberg, 2012). Resulting SAM files were converted to BAM format, sorted, and indexed through samtools (Danecek *et al*., 2021). 50bp tile signal was calculated through the bamCoverage tool in deepTools v3.5.0 suite (Ramírez *et al*., 2016), using bedtools closest (Quinlan and Hall, 2010) to extract the tiles corresponding to the 1kb region centered on each HSF TSS and TTS. MOA-seq data processing was performed as described above before applying the same bamCoverage and bedtools approach to extract tile-based MOA signal.

RNA-seq and MOA-seq datasets have been deposited at the NCBI Short Read Archive, accession number pending.

## Acknowledgements

We thank Peter Hermanson for assistance with plant growth, sampling and library preparation. We thank the Minnesota Supercomputing Institute at the University of Minnesota (http://www.msi.umn.edu) for providing resources that contributed to the research results reported within this article. We thank Daniel Voytas for sharing the Setaria viridis A10 seeds used in this study.

## Tables

**Table S1**. Maize and *Setaria* Gene ID cross-reference table.

**Table S2**. Full DEG table of SvA10 Heat RNA-seq.

**Table S3**. All enriched Gene Ontology terms in *Setaria* heat up-regulated DEGs and shared up-regulated DEGs in 1:1 orthologs of maize and *Setaria*.

**Table S4**. *Setaria* and maize 1:1 ortholog cross-reference table.

## Supplementary Figure Legends

**Figure S1. Multiple sequence alignment of the DNA Binding Domain of the maize and *Setaria* HSFs.** Amino acids are shaded by similarity, with black boxes marking identical and gray boxes marking similar amino acids.

**Figure S2. Dotplot alignments of the two identified *Setaria* tandem duplications.** Transcript annotations and 1kb flanking sequence was extracted for (A) the Sevir.4G250200/Sevir.4G250400 HSFA6a pair and (B) the Sevir.1G026200/Sevir.1G026300 HSFC2a/HSFC2c pair. Gene diagrams along axes indicate gene IDs, transcript annotation (green shading) and a putative HSF TFBS (red shading). The red line in (B) is a result of the proximity of the two annotations to each other, with the downstream ~1kb of Sevir.1G026200 completely overlapping with the upstream ~1kb of Sevir.1G026300. IGV panels depicting the distance between putative tandem duplications are shown for (C) the Sevir.4G250200/Sevir.4G250400 HSFA6a pair and (D) the Sevir.1G026200/Sevir.1G026300 HSFC2a/HSFC2c pair.

**Figure S3. Enriched Gene Ontology (GO) categories in genes up-regulated in response to heat stress.** Enriched GO terms were identified through hypergeometric enrichment analyses in two gene sets, including (A) all genes significantly upregulated in *Setaria*, and (B) all 1:1 orthologs between *Setaria* and maize that were significantly upregulated in both species.

**Figure S4. Expression levels of HSF genes.** (A) *Setaria* and (B) maize HSF expression levels are reported as Counts per Million (CPM) in both control and heat stressed conditions. Expression category and HSF gene IDs are listed above each pair of boxplots.

**Figure S5. Expression heatmap of retained m1:m2 HSF paralogs in maize.** Data includes expression across 247 RNAseq datasets (Zhou *et al*., 2019). Paralog pairs are grouped together, with subclasses denoted on the left and gene IDs denoted on the right. Datasets were organized by hierarchical clustering.

**Figure S6. Chromatin accessibility and modifications over the 1kb region centered on the TSS and TTS of all *Setaria* HSFs.** (A) ATAC-seq coverage, (B) H3K36 trimethylation ChIP-seq coverage, and (C) MOA-seq coverage are presented as heatmaps, with all *Setaria* HSFs split by expression category.

**Figure S7. Chromatin accessibility and modifications over the 1kb region centered on the TSS and TTS of all maize HSFs.** (A) ATAC-seq coverage, (B) H3K36 trimethylation ChIP-seq coverage, and (C) MOA-seq coverage are presented as heatmaps, with all maize HSFs split by expression category.

**Figure S8. Sequence comparison and chromatin information of *Setaria* HSF tandem duplicate pairs.** (A and C) Dotplots and (B and D) heatmaps of the 1kb region centered on the TSS of (A and B) the *HSFA6a* pair *Sevir.4G250200/Sevir.4G250400* and (C and D) the *HSFC2a/HSFC2c* pair *Sevir.1G026200/Sevir.1G026300*. Both dot plots are shaded to highlight key features, including transcript annotations in green and a pair of putative HSF binding sites in red.

